# Closed-loop control of *in vitro* neuronal activity using reinforcement learning after in silico pre-training

**DOI:** 10.64898/2026.07.13.738298

**Authors:** Eduardo Carvalho, José Mateus, Ricardo Pinto, Miguel Aroso, Paulo Aguiar

## Abstract

Controlling specific neuronal dynamics with electrical stimulation is critical for therapeutic neuromodulation, yet deriving optimal control policies remains challenging due to the complex and non-stationary nature of biological neuronal networks. While reinforcement learning (RL) offers a powerful closed-loop control framework, its reliance on prolonged stimulus-driven exploration is difficult to reconcile with the physiological limits of living tissue. Here, we demonstrate an *in silico*-to-*in vitro* transfer strategy that achieves efficient state-dependent control of network bursting in cultured neurons. The transferred policy outperforms heuristic controls, while maintaining constrained stimulation usage. Concurrent calcium imaging reveals the mechanistic basis of the learned policy: the agent optimizes stimulation spatially and temporally, exploiting local network topology and intrinsic physiological temporal dynamics. These results establish *in vitro* brain-on-chip cultures as a tractable stepping stone for RL-based neuromodulation and demonstrate that effective control policies can be derived in biophysically calibrated digital twins and transferred directly to living networks.

## Introduction

The quest to precisely control neuronal activity is an enduring challenge of modern neuroscience, with implications for understanding brain function and developing next-generation therapeutic interventions like deep brain stimulation and brain-machine interfaces^1,2^. Effective neuronal modulation can be pursued using either open-loop or closed-loop control strategies. Open-loop controllers operate on fixed-parameter protocols, delivering stimuli without regard to the system’s ongoing state. While simple to implement, this strategy is fundamentally limited when interacting with a complex, adaptive system like the brain, whose response to input is state-dependent^3,4^. In contrast, closed-loop (or feedback) controllers dynamically adjust their stimulation parameters based on real-time readouts of neuronal activity. Theoretically, such state-dependency offers the potential for more robust, efficient, and personalized neuromodulation^1,5^.

However, translating this theoretical promise into practice in biological neuronal networks (BNNs) presents a formidable challenge. BNNs are not static circuits; they are seemingly stochastic (identical inputs can produce divergent responses) and non-stationary, due to ongoing changes in synaptic connectivity^6,7^. While this set of behaviours is essential for their computational power, it also means that traditional control methods are ineffective, since they are often designed for systems with fixed, predictable dynamics and struggle with large action spaces and non-linear relationships^8^. To mitigate the dimensionality challenge, some closed-loop pipelines restrict the action space *a priori*: they use initial observation periods to identify and select read-out and stimulating electrodes^9–12^. Although this approach has achieved some success, it is limited by the fact that the initial electrode selection can become outdated due to the plasticity and non-stationarity of BNNs^6,13^.

This translation challenge has sparked interest in more flexible control strategies. Reinforcement learning (RL), a machine learning framework for solving sequential decision-making problems^14^, offers a particularly promising and scalable solution. An RL agent can learn an optimal closed-loop policy - that is, a set of rules that determines the best action to take based on the current observed state - through repeated interaction with an environment, without requiring an explicit system model. This makes RL a conceptually attractive framework for adaptive neuromodulation. Yet, a critical gap persists: while RL-based controllers have achieved encouraging results *in silico* within models designed to emulate *in vivo* circuitry^15–18^, their application to actual biological systems has yet to be demonstrated.

A possible step toward closing this gap is to validate RL agents on living neuronal systems that are complex enough to be methodologically relevant, yet simple enough to be experimentally tractable. *In vitro* cultures have historically played this role^19,20^. Under appropriate conditions, these cultures reproduce key features of *in vivo* dynamics - including nonlinearity, adaptability, and non-stationarity - and can exhibit both physiological and pathological activity patterns at the population level. This dynamical fidelity, combined with experimental and ethical accessibility, makes them a practical target for closed-loop control studies^6,21^. In particular, the ability to reliably elicit and disrupt synchronised network dynamics *in vitro* can be directly mapped onto two clinically relevant control objectives: initiating coordinated burst activity, as required when promoting thalamocortical spindles for sleep-dependent memory consolidation^22^, and suppressing pathological synchrony, as pursued in adaptive deep brain stimulation targeting beta oscillations in Parkinson’s disease^1^. Despite this translational potential, the use of RL in BNNs has occurred almost exclusively *in vitro* - and even within this domain, the framework remains largely unexplored. Except for few studies implementing formal RL schemes^23,24^, existing approaches have predominantly relied on stateless parameter optimization (such as adapting electrode selection probabilities based on past performance)^25,26^ rather than learning state-dependent control policies.

These limitations in existing RL-based approaches are not incidental - training RL controllers directly in BNNs poses practical challenges. Defining meaningful state representations from high-dimensional neuronal activity is itself non-trivial^24^; and once training begins, learning reliable control policies requires long training sessions with hundreds to thousands of stimulus repetitions^25,27^. Importantly, prolonged interaction can compromise cell viability, induce stimulus-specific adaptation^10,28^ and drive network drift^29,30^. Together, these factors produce a non-stationary environment where the optimal control policy becomes a “moving target”, progressively devaluing data collected earlier in training.

Therefore, the critical challenge is not merely to bring RL to BNNs, but to do so in a way that is compatible with their physiological constraints. To this end, we present a hybrid pipeline in which an RL agent is first trained *in silico*, using biophysically detailed models calibrated to reproduce the emergent dynamical properties and biological variability of *in vitro* cultures. This training phase allows the agent to explore the policy space under statistically stationary and reproducible conditions before being transferred to living networks. We then demonstrate that the resulting state-dependent control policy generalises to *in vitro* BNNs, successfully eliciting network bursts on demand. Finally, calcium imaging of a subset of experiments reveals that the agent exploits the spatiotemporal dynamics and topological organisation of the underlying network, offering mechanistic insight into the drivers of closed-loop control.

## Results

### Biophysically detailed *in silico* model reproduces key activity dynamics of *in vitro* cultures

We developed a biophysically detailed computational model to recreate the activity of primary hippocampal neuronal networks (Fig. 1a). The model comprised 200 single-compartment Hodgkin-Huxley (HH) neurons, with a composition of 95% pyramidal neurons and 5% parvalbumin-positive (PV+) interneurons. Each neuron featured voltage-gated sodium and potassium channels, along with leak currents representing the intrinsic membrane permeability. Neurons were randomly spatially distributed at a density of 35,000 cells/cm², matching experimental conditions. Synaptic connectivity was implemented with sparse, probabilistic connections, incorporating AMPA, NMDA, and GABA_A_ receptor dynamics, and including short-term plasticity (STP) mechanisms, namely depression and facilitation. To enable direct comparison with experimental recordings, we simulated “virtual” electrodes that captured extracellular spiking activity. Individual neuron and synapse parameters were fixed based on literature-derived experimental data, whereas the network’s connectivity parameters were tuned to reproduce the characteristic dynamics of our *in vitro* preparations - embryonic (E18) rat hippocampal neurons cultured on microelectrode array (MEA) chips (Fig. 1b).

**Figure 1.**
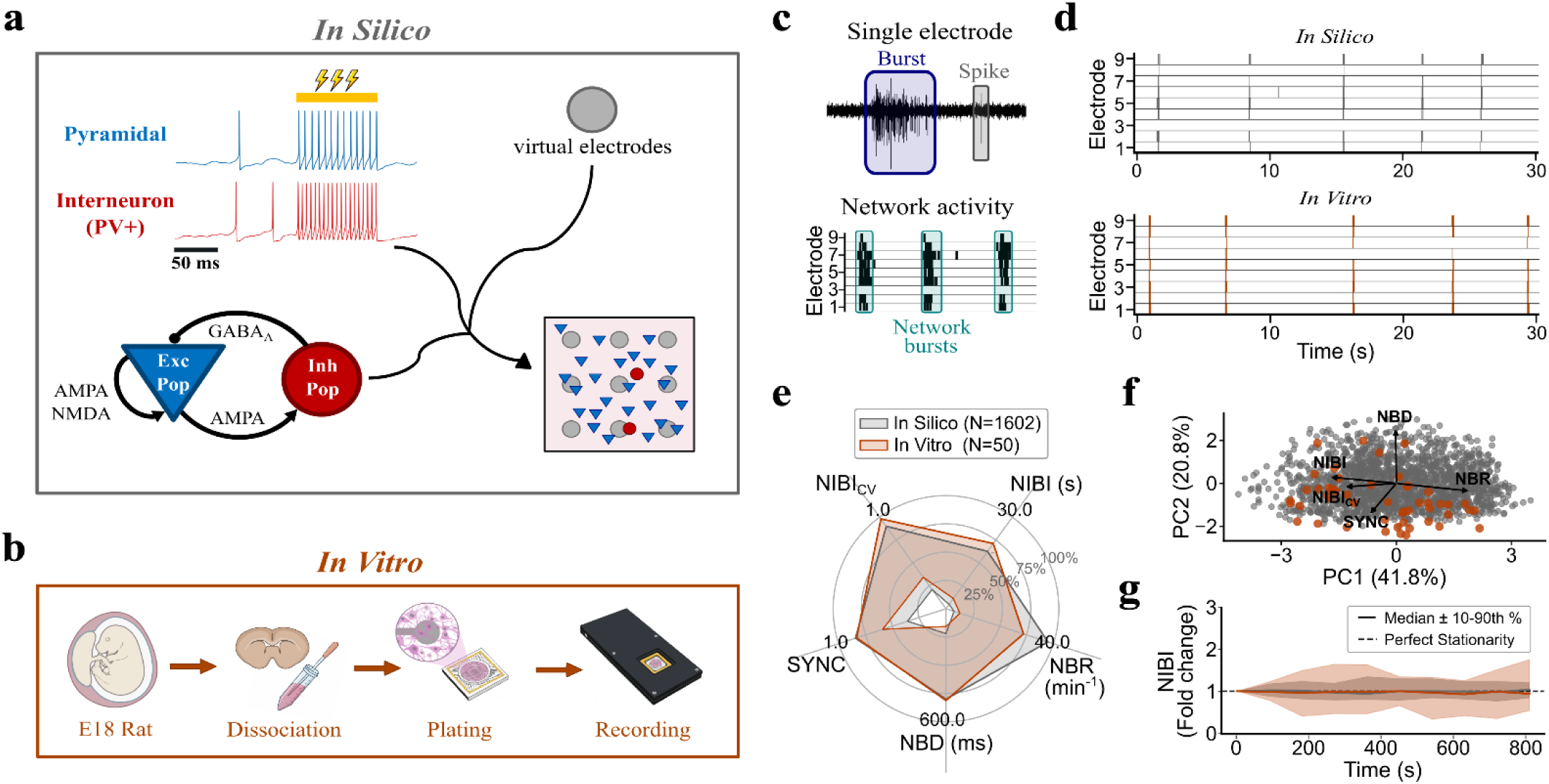
In silico recapitulation of the bursting phenotype observed in in vitro hippocampal networks. **(a)** Schematic of the biophysical in silico model consisting of 200 Hodgkin-Huxley (HH)-type neurons, containing voltage dependent sodium and potassium currents, sparsely connected via AMPA, GABA_A_ and NMDA receptors, using virtual electrodes to measure network activity. The cell placement follows a Poisson process with density 35,000 cells/cm^2^. The pyramidal and interneuron populations are represented by blue triangles and red circles, respectively. Electrodes are grey circles. **(b)** Protocol for in vitro cell culture. Hippocampal cells were obtained from embryonic day 18 (E18) rats. Activity was recorded between DIV 17–42. **(c)** Electrophysiology schematic showing simulated single-channel burst/spike detection and network-wide bursting. **(d)** Raster plots illustrating the activity of in silico networks (top) and in vitro cultures (bottom). **(e)** Quantification of network activity metrics. Filled area represents the 2.5^th^ to 97.5^th^ percentiles for each metric. (NBR: network burst rate; NIBI: network inter-burst interval; NIBI_CV_: coefficient of variation of NIBI; SYNC: synchrony; NBD: network burst duration). **(f)** PCA projection of individual network dynamics, showing that NBR and NIBI are anti-correlated and nearly orthogonal to NBD. **(g)** Fold change of NIBI of 15-minute baseline recordings for both in silico (N=50, grey) and in vitro (N=20, salmon) networks. Shaded areas indicate the 10^th^ to 90^th^ percentiles

*In vitro* cultures showed highly reproducible and well-established activity patterns across preparations, consisting of individual spikes and bursts (brief periods of high-frequency firing) that spontaneously organized into network bursts (NBs) detectable across multiple electrodes^7,31^. Our *in silico* model successfully recapitulated these spatiotemporal dynamics, with NBs emerging spontaneously across the array of “virtual” electrodes (Fig. 1c). Notably, the *in silico* networks exhibited significantly less non-stationarity than the *in vitro* cultures, maintaining stable network inter-burst interval values over time (Fig. 1g; similar results for other metrics are not shown). Such relative stability is anticipated; although the model incorporates STP to capture rapid firing dynamics, the underlying structural connectivity and synaptic weights remain fixed, providing a stationary baseline compared to the stochastic drift of living tissue.

To quantitatively validate the model, we used five metrics: network burst rate (NBR), network burst duration (NBD), network inter-burst interval (NIBI) and its coefficient of variation (NIBI_CV_), and electrode synchrony (SYNC). All simulated metrics fell within experimentally observed ranges (Fig. 1e), demonstrating that the model accurately recapitulates the main dynamics of *in vitro* network activity. Principal component analysis (PCA) further revealed that the model captured not only parameter ranges but also the electrophysiological phenotype (Fig. 1f), effectively serving as a functional digital twin of the *in vitro* networks.

### Reinforcement learning agent generalizes network burst control in silico

After the calibration of the simulated model, we designed a reinforcement learning pipeline to maximize the immediate probability of eliciting a network burst (NB) while minimizing stimulation usage. This objective was chosen because NBs represent well-defined, temporally localized events of maximal network coordination, providing a natural and explicit reward signal, and a stringent test of whether the agent has learned to exploit the network’s internal excitability state. This framework for state-dependent control was further motivated by clinical contexts in which targeted promotion of synchronised activity is desirable^22^.

Reliable generalisation across network simulations required grounding the agent’s decisions in observable electrophysiological features that reflect excitability in a simulation-invariant manner. Stimulation efficacy depends nonlinearly on both temporal context (e.g., recovery from previous NB) and spatial context (e.g., electrode-specific network coupling). We therefore designed a state representation integrating both dimensions (see *Methods: State representation*). Each episode consisted of sequential 200 ms stimulation steps (Fig. 2a). Episodes terminated upon NB detection and resumed only after a 200 ms quiescent period, ensuring the end of the previous NB.

**Figure 2.**
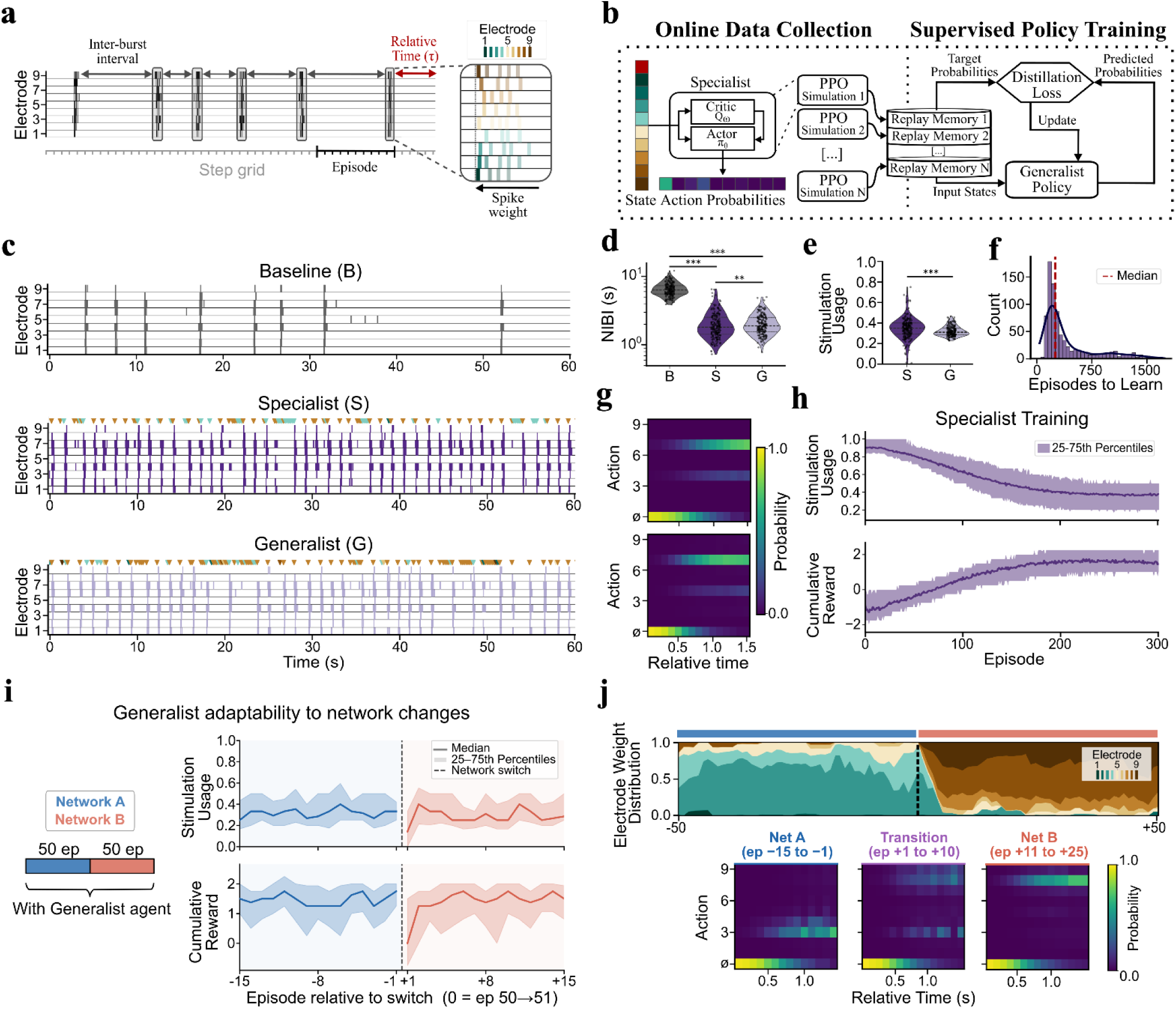
Design and *in silico* validation of the *Generalist* policy. **(a)** Schematic depicting how the state representation includes past recent history. The temporal feature (in red) is derived by dividing the elapsed time since the last network burst (NB) by the median of the last five network inter-burst intervals (NIBIs). Spatial features (electrode weights) are calculated by summing spike weights per electrode over the preceding five NBs. **(b)** Policy distillation pipeline. State-action pairs obtained during *Specialist* agent training are aggregated to train a single *Generalist* agent. **(c)** Representative raster plots of network activity under *Baseline*, *Specialist*, and *Generalist* policies. Coloured triangles indicate stimulated electrodes (colours correspond to electrode in **a**). **(d,e)** Validation on held-out simulations (N=200). **(d)** Average NIBI comparison. **(e)** Stimulation usage (fraction of steps with stimulation). Note that while the performance difference is statistically significant, the median difference in NIBI (66 ms) is below the simulation time-step (200 ms), indicating practical equivalence. **(f)** Distribution of training episodes required for *Specialist* policy convergence (N=999; median=248 episodes) **(g)** Stimulation probability heatmaps as a function of normalized time since the last NB for *Specialist* (top) and *Generalist* (bottom) policies for the network in panel **c**. **(h)** Learning trajectories (cumulative reward and stimulation usage per episode) for all *Specialist* agents during training. Line indicates median and shaded areas the 25^th^-75^th^ percentiles. **(i)** Generalist agent performance during a network-switch stress test, in which the same agent was exposed sequentially to two distinct simulated networks (Network A and Network B), each for 50 episodes. Top: stimulation usage per episode. Bottom: cumulative reward per episode. Line indicates median and shaded areas the 25th– 75th percentiles. The dashed vertical line indicates the network switch (episode 50→51). **(j)** Example of electrode weight distribution and action probability heatmaps across the network transition. Top: evolution of per-electrode state weights over the 50 episodes preceding and following the switch (dashed vertical line). Bottom: stimulation probability heatmaps as a function of normalised time since the last NB, averaged over three epochs - late Network A (episodes −15 to −1), transition (episodes +1 to +10), and late Network B (episodes +11 to +25). *Significance levels are denoted by * (p<0.05), ** (p<0.01), and *** (p<0.001)*.

To achieve robust performance across different networks, we used a two-stage training strategy (Fig. 2b). First, we turned to online learning, where individual agents were trained from scratch using Proximal Policy Optimization^32^ (PPO) on multiple simulated networks, generating environment-specific (hereafter termed *Specialist*) stimulation policies. *Specialists* showed rapid learning progression, with cumulative rewards and stimulation usage (fraction of actions where stimulation was used) reaching asymptotic performance within 250 training episodes (Fig. 2f,h). The state-action distributions from these agents were aggregated for policy distillation (Fig. 2b). Then, an agent was trained on these state-probability pairs to produce a *Generalist* policy that minimized the Kullback-Leibler (KL) divergence of the outputted probabilities across all training environments.

We then tested the *Generalist* policy on a subset of simulated networks (excluded from policy distillation) and compared its performance to the respective *Specialists*. Consistent with the premise that fundamental excitability dynamics are conserved across networks, the *Generalist* agent’s action probabilities often matched those of network-specific *Specialist*s (Fig. 2g). Both the *Specialist* and *Generalist* agents significantly decreased the NIBI relative to baseline in most networks (Fig. 2d), though the degree of improvement varied substantially across networks. Notably, while maintaining comparable NIBI reduction capability to *Specialists*, the *Generalist* policy consistently used less stimulation (Fig. 2e). This combination of preserved efficacy and reduced behavioural variability is a hallmark of successful policy generalization^33^. As a stress test of the *Generalist* policy’s adaptability, we exposed the same agent sequentially to two distinct simulated networks (Network A and Network B), each for 50 episodes, representing an abrupt and complete change of network context (Fig. 2i). Following the switch, cumulative reward and stimulation usage transiently shifted before restabilising within 5 episodes (consistent with the time required for the state representation to be computed exclusively from Network B history), demonstrating rapid online adaptation to a new network context.

To understand the mechanism underlying this rapid adaptation, we examined how the agent’s action preferences evolved across the transition (Fig. 2j). The electrode weight distribution - a key component of the state representation - shifted markedly at the switch point, with previously dominant electrodes losing weight in favour of those more informative for Network B. Correspondingly, the action probability heatmaps show that the policy redistributed stimulation probability across a different electrode set within the first episodes following the switch, before converging to a stable Network B strategy by episodes +11 to +25. These results indicate that the Generalist policy does not rely on a fixed stimulation strategy but instead continuously adapts its actions to the evolving state representation, tracking changes in network excitability and electrode-specific coupling without any explicit network identification step.

### Online learning induces network fatigue *in vitro* without improving control efficacy over pre-trained *Generalist* agent

To compare our approach against standard online learning, we first assessed the feasibility of training *Specialist* agents from scratch directly on *in vitro* networks (Protocol 1, Fig. 3a). While technically achievable, online learning highlighted an inherent challenge in biological RL: the balance between policy convergence and network stability. Stabilizing an effective policy typically required a comparable number of episodes to *in silico* training (Fig. 3b), corresponding to approximately five minutes of stimulation (Fig. 3c). However, this training period was already sufficient to trigger a measurable degradation in network activity, which we henceforth define as “network fatigue”. Specifically, most networks reduced spontaneous burst frequency during the final baseline (Fig. 3h,i). This fatigue caused a confounding order effect: *Generalist* policies tested after a *Specialist* training phase showed longer network inter-burst intervals (NIBIs) compared to *Generalist* policies tested first (Fig. 3g). An alternative comparison between online learning and our proposed approach would likely require a reduction in stimulation density to minimize network fatigue - an adjustment that would, in turn, significantly extend the time required to reach policy convergence.

**Figure 3.**
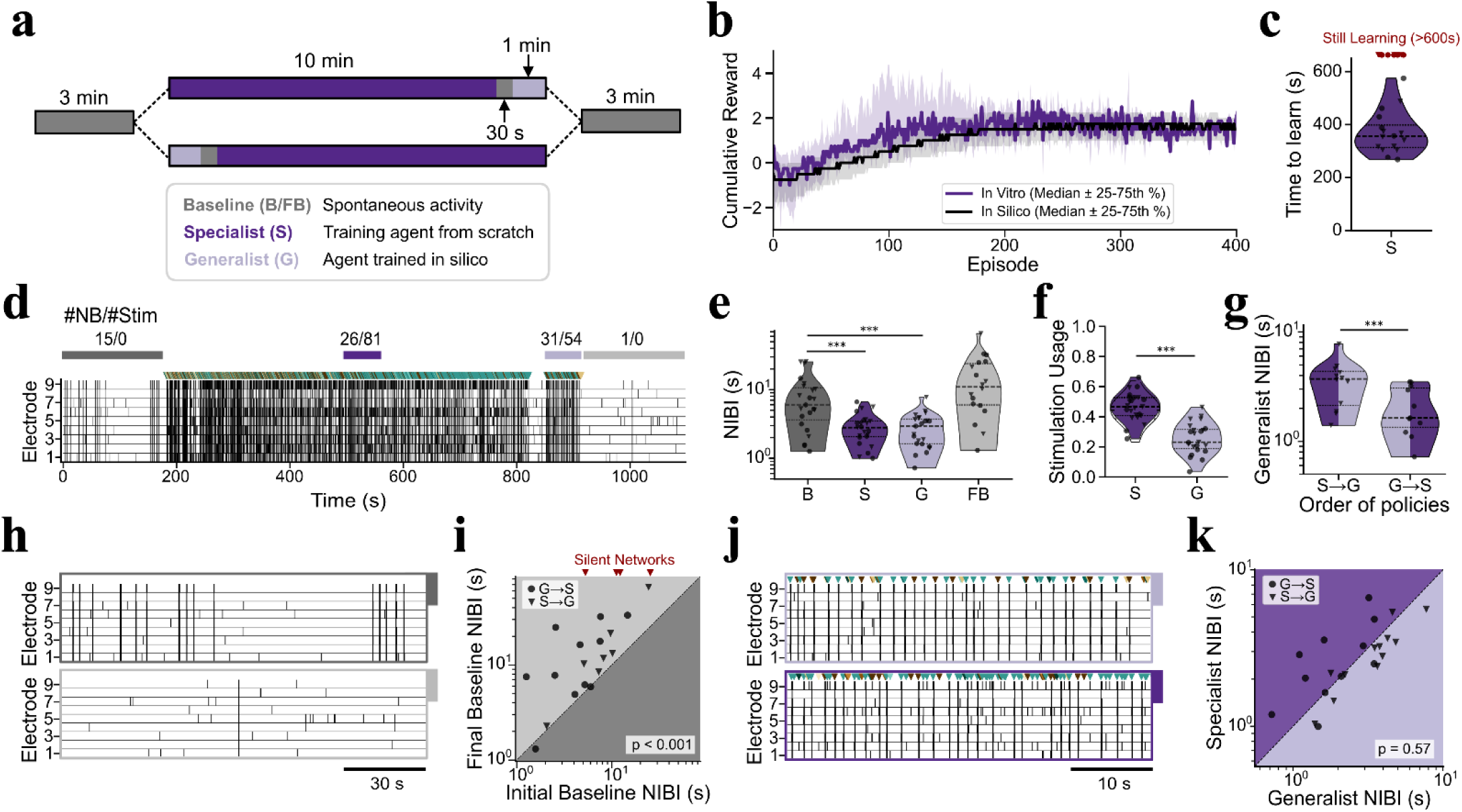
Benchmarking the *Generalist* policy against naive online learning. **(a)** Schematic of the experimental protocol. Two testing sequences were used to control for order effects between *Generalist* and *Specialist* agents**. (b)** Learning trajectories (cumulative reward per episode) of in vitro *Specialist* agents compared to in silico counterparts. Shaded areas indicate the 25^th^-75^th^ percentiles. **(c)** Distribution of time until cumulative reward convergence for *Specialist* agents. About 70% (16/23) converged to a policy within the 10-minute window. **(d)** Representative raster plot of a full experiment exhibiting significant network fatigue. Coloured bars indicate time windows detailed in **h** and **j**; the number of network bursts (NB) and stimuli for each window are indicated above. **(e)(f)** Comparison of control efficacy (NIBI, **e**) and efficiency (Stimulation usage, **f**) across conditions (N=23). **(g)** Impact of experimental order on *Generalist* agent performance (NIBI). Groups: S→G (N=12) vs. G→S (N=11). **(h)** Detailed view of spontaneous activity during Initial and Final baselines from the raster in **d**, of a network with marked network fatigue. **(i)** Quantification of network fatigue. Scatter plot compares spontaneous mean network inter-burst interval (NIBI) before (Initial) and after (Final) policy deployment. Silenced networks (defined as having <2 NBs in the Final Baseline) account for 22% (5/23) of experiments. **(j)** Detailed view of network activity under *Generalist* and *Specialist* control from the raster in **d**. **(k)** Pairwise comparison of control efficacy (NIBI) between *Specialist* and *Generalist* policies. The *Generalist* agent matches *Specialist* performance without online training in the biological preparation. *Significance levels are denoted by * (p<0.05), ** (p<0.01), and *** (p<0.001)*.

We also noted that initial training trajectories deviated from *in silico* predictions during early training phases (Fig. 3b). Specifically, we observed an initial increase in median cumulative reward driven by less responsive networks; in these cases, extended periods of network quiescence allowed the agent to accumulate positive rewards for withholding stimulation, thereby inflating the cumulative signal prior to policy convergence (see *Methods: Reward function*). Expectedly, these networks were also significantly more prone to failing convergence criteria.

Regarding task performance, the pre-trained *Generalist* policy proved robust against the *Specialist* benchmark. When evaluated during the converged performance phase (or the final minute of training for non-converging networks), the *Specialist* agents yielded no significant improvement in NIBI reduction over the *Generalist* agent (Fig. 3e,k). In fact, the *Generalist* agent demonstrated greater parsimony, achieving comparable control with significantly less stimulation (Fig. 3f) - recapitulating what was observed during *in silico* training (Fig. 2e,h).

### The pre-trained *Generalist* agent outperforms heuristic controls in evoking *in vitro* network bursts

Despite the clear advantages of using a pre-trained *Generalist* agent over naive online learning, we still sought to determine if it was approaching the maximum possible performance of our control task. To this end, we compared the *Generalist* against two heuristic controls (Protocol 2, Fig. 4a), each deliberately depriving a key aspect of the agent’s decision-making. The “*Without Best*” policy restricted stimulation to all electrodes except the one empirically identified as highest yielding for that network, removing the agent’s ability to exploit its single most successful action. The “*Random*” policy applied spatially randomized stimulation at the same overall stimulation rate (stims/min) as the *Generalist*, removing any learned spatial selectivity while preserving the total stimulation budget. Together, these controls probe the *Generalist* from two directions: whether its advantage stems from knowing which electrode to stimulate, or from the overall rate of stimulation.

**Figure 4.**
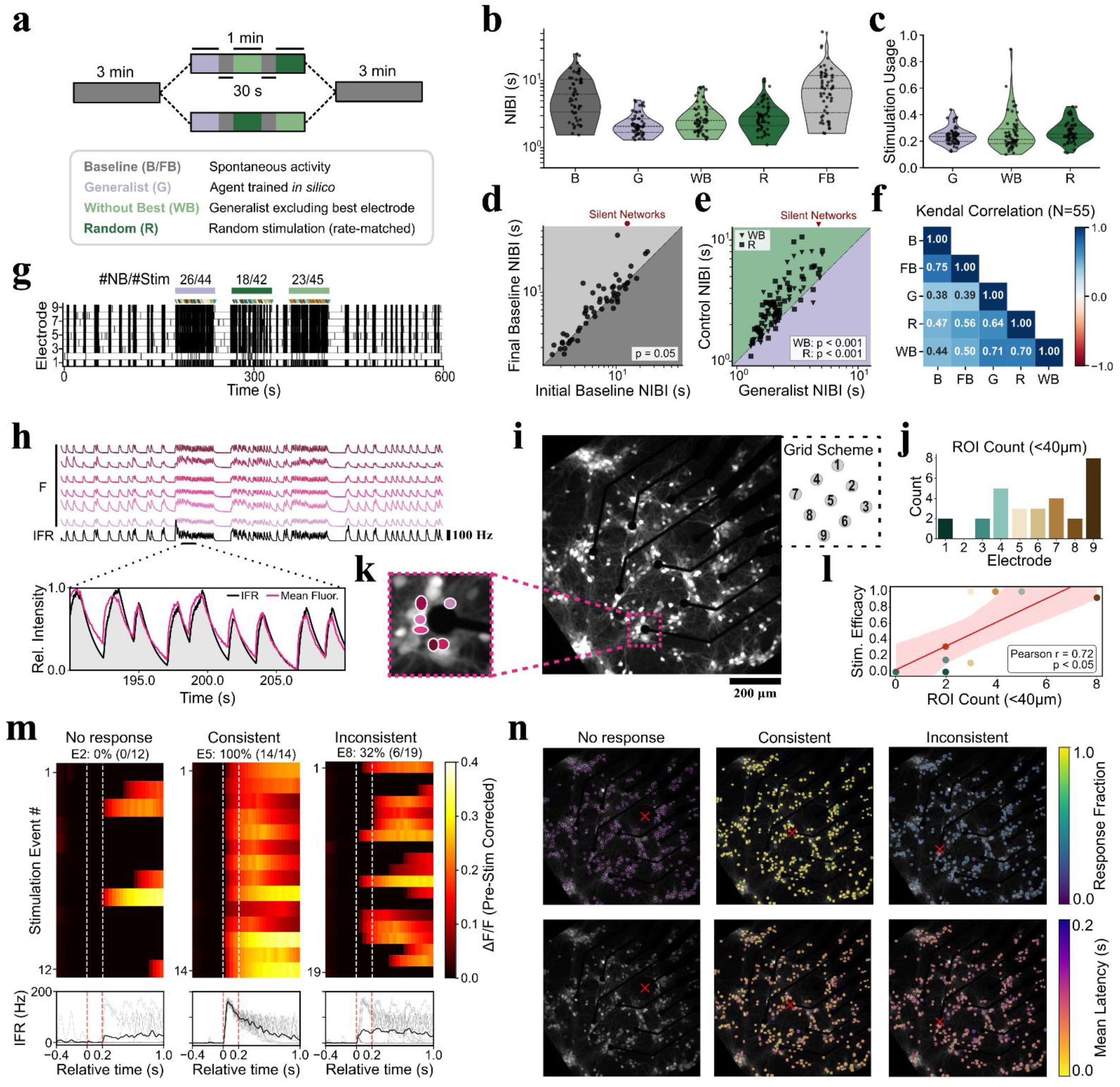
The Generalist reinforcement learning agent outperforms heuristic controls by exploiting local network topology and temporal dynamics. **(a)** Schematic of the experimental protocol. Two testing sequences were used to control for order effects between Without Best and Random policies**. (b)(c)** Comparison of control efficacy (NIBI, **b**) and efficiency (Stimulation usage, **c**) across the tested policies. **(d)** Quantification of network fatigue. Scatter plot compares spontaneous mean network inter-burst interval (NIBI) before and after deployment of the stimulation policies. **(e)** Pairwise comparison of control efficacy (NIBI) between Generalist and control policies. Most control policies perform significantly worse than their Generalist counterparts. **(f)** Kendall rank correlation matrix of NIBI across all phases (N=55). High correlation between initial (B) and final baselines (FB) indicates temporal stability of intrinsic network dynamics, while moderate correlations between baseline and active stimulation policies (G, R, WB) reflect stimulation-induced modulation of network ranks. **(g)** Representative raster plot of one full experiment where concurrent calcium imaging was performed. Coloured bars denote the sequence of applied stimulation policies, and the numbers above indicate the total count of network bursts within each phase. **(h)** Instantaneous firing rate (IFR) of Electrode 9 and calcium fluorescence (F) traces from neighbouring ROIs (highlighted in **k**). The top panel shows traces across the full experiment. The bottom panel shows a magnified epoch during Generalist agent testing, with both signals locally min-max normalized [0, 1] to highlight their shape correlation. **(i)** Representative calcium fluorescence (jGCaMP8m) image of the neuronal network cultured on the microelectrode array at DIV18. The inset (top right) details the grid scheme and numbering of the 9 electrodes. The dashed magenta box indicates the magnified region shown in **k**. Scale bar, 200 µm. **(j)** Quantification of active somatic ROIs detected within a 40 µm radius of each recording electrode. **(k)** Magnified view of the region highlighted in **i**, detailing the identification of individual active somatic ROIs (magenta outlines) immediately adjacent to the electrode. **(l)** Scatter plot showing the Pearson correlation (r = 0.72, p < 0.05) between electrode efficacy at eliciting NBs and the local ROI count (<40 µm). The red line indicates the linear trend with its corresponding confidence interval (shaded). **(m)** Pulse-by-pulse responses for three representative electrodes demonstrating “No response” (Electrode 2), “Consistent” (Electrode 5), and “Inconsistent” (Electrode 8) activation profiles. Top: Heatmaps showing the pre-stimulus-corrected local calcium fluorescence (ΔF/F) for individual stimulation events. Bottom: Corresponding network IFR traces for all individual network responses (grey) and the response average (black). Percentages denote the fraction of stimulation events that elicited a detectable response within the evaluation window (red dashed lines). **(n)** Spatial maps of network recruitment following stimulation of the three representative electrodes detailed in **m** (stimulation sites marked by red crosses). Top row: The fraction of trials in which each individual ROI exhibited a significant calcium response, visualizing the spatial extent and reliability of network activation. Bottom row: The mean response latency in seconds for each ROI across successful actions, illustrating the temporal propagation of evoked activity.

All stimulation policies successfully reduced the mean network inter-burst interval (NIBI) relative to spontaneous baseline activity. However, the *Generalist* agent achieved greatest reduction in NIBI (Fig. 4b) while maintaining comparable stimulation usage (Fig. 4c). Pairwise comparisons further underscored this superiority, demonstrating that the *Generalist* agent significantly outperformed its heuristic counterparts on an experiment-by-experiment basis (Fig. 4e).

We next investigated whether the “network fatigue” observed during naive online learning (Protocol 1) persisted under this shorter evaluation scheme. While Protocol 2 showed a trend toward post-stimulation increase in NIBI (p = 0.052), the magnitude of this effect was drastically reduced compared to Protocol 1. Specifically, Protocol 1 induced severe exhaustion after 11 minutes of active stimulation, whereas Protocol 2 produced only mild drift after 3 minutes (d = 1.04 vs. d = 0.39). This confirms that sustained, high-dose stimulation is the primary driver of network fatigue. To ensure this mild drift did not systematically bias the evaluation of the subsequent heuristic controls, we conducted a validation protocol consisting of three consecutive 1 minute-phases of the *Generalist* policy. This validation revealed no significant degradation in performance over time (p = 0.31, Fig. S1). Furthermore, we confirmed neither *Random* (p = 0.81) nor *Without Best* (p = 0.81) NIBI distributions differed significantly by deployment order.

A striking feature of these cultures is the sheer variance in their spontaneous baseline activity, which spans a full order of magnitude (hence the log-scale representation in the NIBI plots). Such variability poses a natural question: do these differences in spontaneous NIBI statistics dictate how excitable the networks are, or can the *Generalist* agent simply overpower them? To answer this, we first correlated each network’s spontaneous baseline NIBI with its performance under each policy (Fig. 4f) and found consistent moderate positive associations across all three (τ = 0.38–0.47): networks with shorter spontaneous inter-burst intervals tended to achieve shorter intervals under closed-loop control as well. We also compared the performance ranks of the different stimulation policies across individual networks and observed similarly strong cross-policy correlations (τ = 0.64–0.71), indicating that no policy can substantially reorder networks relative to the others. Taken together, these results suggest that network identity (rather than policy choice) is the dominant determinant of NB elicitation performance, with each preparation’s intrinsic properties setting an effective ceiling on NIBI reduction. This ceiling likely reflects a combination of the network’s spontaneous refractory-like dynamics and preparation-specific factors such as electrode-network coupling via the MEA interface.

### Spatiotemporal calcium dynamics reveal the topological drivers of electrode-specific stimulation efficacy

To investigate the spatial network properties driving the *Generalist* agent’s choice of electrodes and its subsequent performance, we performed concurrent calcium imaging during a subset (N=5) of the Protocol 2 evaluation experiments. We first examined the relationship between the instantaneous firing rate (IFR) derived from electrode spikes and the calcium fluorescence traces (*F*) of nearby segmented neuronal somata (ROIs) (Fig. 4h,k). Across the imaging dataset, local calcium dynamics exhibited a strong positive correlation with the underlying electrophysiology. Notably, this high correlation was maintained even when only a single active ROI was detected within a 40 µm radius of the recording electrode. This indicates that the presence of at least one optically active soma within this immediate neighbourhood is a reliable proxy for local spiking.

Building on this electro-optical coupling, we next assessed how this local cellular density dictated the efficacy of targeted electrical stimulation. Pulse-by-pulse analysis of individual electrodes revealed a stark dependency on local ROI count (Fig. 4m). For instance, stimulation of Electrode 2, which lacked any detectable adjacent ROIs, completely failed to evoke NBs (0% response rate). Conversely, stimulation of Electrode 5, which was surrounded by three active ROIs, consistently recruited the broader network, with a 100% success rate. Intermediate cases, such as Electrode 8 with two adjacent ROIs, yielded inconsistent network recruitment (32% response rate). Notably, this particular network had four distinct electrodes with stimulation response rates exceeding 90%. With the exception of Electrode 6, these highly efficacious electrodes corresponded directly to the ones with the highest local ROI counts (Fig. 4l). This spatial redundancy explains why the *Without Best* policy achieved a comparable number of NBs relative to the *Generalist* agent (23 vs. 26 NBs), as the algorithm could readily route control through alternative electrodes.

Spatial mapping of these responses further revealed that NBs evoked by the highly reliable Electrode 5 exhibited substantially lower activation latencies across the culture, indicating a more efficient temporal propagation of the perturbation (Fig. 4n). Across some stimulation trials, network recruitment latencies extended up to 100 ms. Crucially, this physiological propagation delay directly validates the 200 ms time-step duration chosen for the reinforcement learning task; it ensures the biological network has sufficient time to respond to the stimulus before the agent evaluates the new state and selects its subsequent action.

While local somatic density was a strong overall predictor of efficacy, we did observe specific exceptions to this rule. We noted two instances across our dataset where an electrode exhibited high spiking activity without any visible adjacent ROIs (Fig. S2); careful inspection of one case revealed that this was caused by active cells situated directly atop the opaque metallic electrode lead, which obscured their fluorescence. Additionally, in two of the five tested networks, we found two moderately to highly efficacious electrodes that lacked visible adjacent ROIs, high local spiking activity, and strong correlations with the network calcium traces (Fig. S2). This suggests that the immediate presence of a soma is not strictly necessary for successful network excitation.

## Discussion

Our *in silico* framework was designed to recapitulate key activity dynamics observed *in vitro* - such as bursting statistics and response probabilities - within a biophysically grounded yet computationally tractable environment. Unlike biological neuronal networks (BNNs), which are characterized by non-stationarity and network drift over extended timescales^24,34^, the simulated environment maintains statistical stationarity. Such stability provides the clean signal required for robust policy learning. It insulates the agent from confounding physiological drift, ensuring that training optimizes for genuine causal dependencies rather than overfitting to transient excitability states. Consequently, the model functions as a standardized benchmark, enabling the rigorous training and validation of agents under reproducible conditions prior to their deployment in BNNs.

We conceptualized the closed-loop neuromodulation task as a nonlinear contextual bandit problem, where the objective is to map an instantaneous network state to the optimal stimulation target that maximizes the immediate probability of a network burst while penalizing stimulation usage. This framing is biologically appropriate for two reasons. First, the control goal is inherently ‘greedy’ - prioritizing immediate responsiveness over long-horizon planning. Second, our data confirms that electrode efficacy is not static but varies nonlinearly with the network’s recovery status (i.e., whether the network is refractory or excitable). However, while the theoretical formulation aligns with the bandit framework, the standard algorithmic solutions are inadequate for real-time implementation. Nonlinear bandit algorithms (e.g., NeuralUCB^35^) typically rely on explicit uncertainty quantification, continuously calculating confidence bounds to guide exploration. This introduces a computational overhead that grows with experimental duration, often exceeding the sub-millisecond latency desirable for precise feedback. Furthermore, these methods may struggle with the non-stationarity of biological tissues^36^. To resolve this conflict between theoretical framing and practical latency, we used an adapted version of the Proximal Policy Optimization (PPO) algorithm. This choice offers two main advantages. First, PPO learns a reactive policy capable of low-latency inference (single forward pass), effectively solving the contextual bandit challenge within the strict timing constraints of the biological loop. Second, unlike specialized bandit algorithms, PPO provides a unified framework that naturally generalizes beyond the greedy setting: it allows us to solve the immediate bandit problem (discount factor *γ* = 0) while retaining the architectural flexibility to incorporate long-horizon credit assignment (*γ* > 0) should the control objectives evolve.

It is also worth noting that the success of this algorithm relies on a state representation explicitly designed to capture instantaneous network excitability. By encoding the recent history of network bursting activity, the state vector provides the agent with a proxy for both recovery status and NB spatial propagation. This design choice is validated by the strong correlation observed between the agent’s learned electrode probabilities and the response efficacy of those electrodes - effectively, the agent learns to ‘trust’ certain electrodes based on the state context. Moreover, the sufficiency of this state representation explains why policy distillation was highly effective. Because the state vector successfully disentangles the noisy biological dynamics, the mapping from *State* to *Optimal Action* becomes relatively stable and distinct. While end-to-end RL could theoretically infer these latent excitability dynamics directly from raw spike trains^37^, such an approach would require orders of magnitude more data. The success of our compact feature set suggests that complex deep-learning architectures yield diminishing returns for this specific class of control problems, where dynamics are dominated by identifiable recovery variables.

Deployment *in vitro* revealed a critical physiological constraint: the cumulative cost of interaction. While Protocol 2 (∼3 minutes of cumulative stimulation) induced only mild network fatigue, Protocol 1 (∼11 minutes of cumulative stimulation) resulted in a pronounced effect. This reduction in spontaneous activity aligns with previous findings that sustained low-frequency stimulation drives suppression of network bursting^28,29,38^. The contrast between our two protocols suggests that post-stimulation fatigue is a function of total stimulation density over time, whereby longer stimulation windows result in a stronger effect. Because of this strict physiological limit, transferring a fully trained agent is not just convenient, but biologically necessary. It allows us to skip the fatigue-inducing phase of RL and apply control while the network is still highly responsive. Crucially, safeguarding the cells in this way does not require a compromise in performance. Our data confirms that *in silico* pre-training was not inferior to online learning.

The superior performance of the pre-trained *Generalist* agent over heuristic controls demonstrates that it learned a spatially optimized stimulation strategy rather than simply driving non-specific excitability. Interestingly, our rank correlation analysis revealed a persistent hierarchy of network responsiveness across all stimulation paradigms. This temporal stability strongly supports the initial assumption of the presence of an intrinsic network refractory period - likely driven by synaptic vesicle depletion and slow afterhyperpolarization currents - that fundamentally bounds the maximum achievable NB rate, regardless of the controller applied. In contrast, the spatial dimension of stimulation efficacy is heavily dependent on the specific topology of the underlying culture. The varying success of the heuristic policies highlights this: the “*Without Best*” policy severely degrades performance in networks possessing only a single highly efficacious stimulation electrode, whereas networks with a distributed availability of “good” electrodes remain highly responsive to both *Without Best* and *Random* strategies.

Concurrent calcium imaging revealed that the *Generalist* agent learned to exploit the spatial embedding of electrodes within the network’s functional topology. We found that the presence of active somatic ROIs within a 40 µm radius of an electrode reliably predicted its ability to evoke network bursts. However, it is important to contextualize this finding within the broader biophysics of extracellular stimulation. While traditional models (including our own) often employ Stoney’s relationship^39^ to define a spherical volume of somatic activation, both computational^40^ and experimental^41^ works suggest that electrical stimulation more readily recruits traversing axonal fibres rather than local somata. In our *in silico* framework, applying Stoney’s assumption to purely somatic compartments was a necessary compromise to balance biological realism with computational tractability. Consequently, our model seems to be underestimating the true density of efficacious stimulation hubs by not accounting for the activation of distant cells via passing axons. Nevertheless, this somatic-centric abstraction aligns perfectly with the physical constraints of our *in vitro* setup. The 30 µm diameter electrodes used in our MEAs are optimized for somatic interfacing and lack the spatial resolution required for the reliable detection or targeted isolation of individual axons^42^.

By shifting the data-intensive burden of policy discovery to an *in silico* environment, our pipeline resolves the fundamental conflict between the sample inefficiency of RL and the experimental limitations of BNNs. While the current implementation demonstrates feasibility, future iterations of the RL scheme could be refined to explicitly incorporate physiological constraints (e.g., long-term depression, stimulus-specific adaptation) or continuous-time dynamics (i.e., action not limited to a fixed time step), thereby minimizing the physiological cost of intervention. Crucially, this framework lays the groundwork for addressing more complex, functionally relevant neuromodulation challenges. By extending it to tasks such as stabilizing pathological states via network desynchronization^43^ or orchestrating dynamics within multi-population networks with controlled architectures^44,45^, we can move beyond simple activity maximization toward the precise regulation of more complex network dynamics. Ultimately, our results support the feasibility of this *in silico*-to-*in vitro* transfer strategy, providing a proof-of-concept for how neuronal interfaces operating under strict data scarcity and safety constraints (such as adaptive DBS) might leverage simulation-based pre-training to accelerate closed-loop control without requiring extensive online learning.

## Methods

### Computational model

The *in silico* model was implemented in Python using the well-established NEURON environment^46^ and consists of several submodules. Neurons, synapses, and network parameter values are based on experimental data and literature, when possible, with the remaining parameters chosen to ensure the simulations reproduce activity seen in primary *in vitro* hippocampal cultures.

#### Neuron models

We used an HH-type neuron model^47^, with rate constant expressions optimized for hippocampal pyramidal neurons^48^:

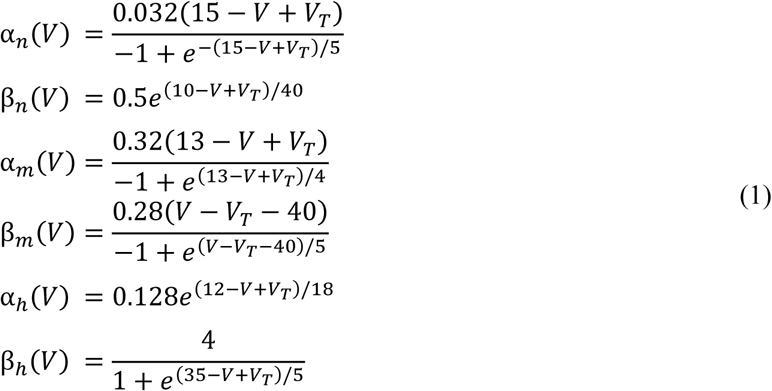

where *V* is the membrane potential of the neuron, and *V_T_* a constant that allows the adjustment of the spiking threshold. Parvalbumin positive (PV+) interneurons were modelled as a variant of the pyramidal neurons, featuring a smaller diameter and distinct conductances, as described in Supp. Table 1.

#### Noise

Independent membrane potential fluctuations were induced in every neuron, using NEURON’s *NetStim* mechanism to generate trains of presynaptic stimuli following a homogeneous Poisson process^49^. The current injected in each neuron is estimated by:

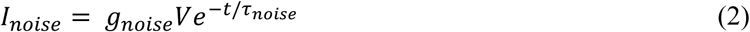

Different noise levels were obtained by changing the stimuli rate 1/*τ_noise_* and the maximum conductance *g_noise_*. Reference values for these constants are described in Supp. Table 1.

#### Synapses and plasticity

The following synaptic receptors were included in the model: AMPA, NMDA and GABA_A_. All synaptic conductances were modelled with biexponential kinetics, governed by rise and decay time constants, denoted as *τ_rise_* and *τ_decay_*, respectively. The maximum conductance of each synapse is defined by *ĝ*. Values for these parameters were taken from Romani et al.^50^ and are shown in Supp. Table 2. The NMDA conductances follow the model described in Aguiar et al.^51^.

For NMDA conductances, a magnesium block was included, modelled by a dimensionless multiplicative factor *f_Mg_*, described by the biophysical model of Jahr and Stevens^52^:

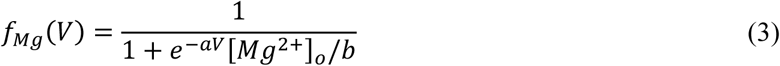

where *V* is the membrane potential of the postsynaptic neuron, [*Mg*^2+^]*_o_* is the extracellular magnesium concentration, which is set to 1 *mM* as in the original model, and *a* and *b* are rate constants, with the original values of 0.062 *mV*^−1^ and 3.57 *mM*, respectively. In addition, from the total nonspecific cationic current passing through the NMDA channels, a fraction of 10% was set as a calcium current^53^.

All synapses were subject to short-term depression and facilitation, modelled using the probabilistic framework proposed by Fuhrmann et al.^54^. This model is described by the following differential equations:

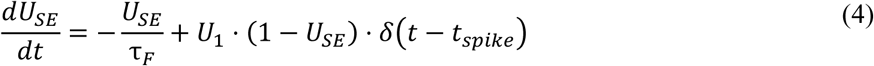

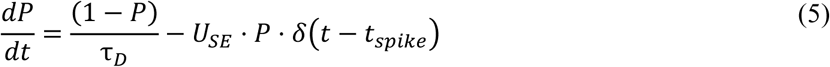

, where *U_SE_* is the amount of resources used in a release at time t, *U*_1_ is a constant determining the step increase in *U_SE_*, *P* is the probability of a vesicle being available for release at time t, *δ*(*t*) is the Dirac delta function, and *t_spike_* is the timing of the presynaptic spike. The values of the parameters are shown in Supp. Table 2.

#### Recording extracellular potentials

The extracellular potentials at each electrode were calculated as the sum of multiple point sources using a modified version of the quasistatic approximation of Maxwell’s equations:

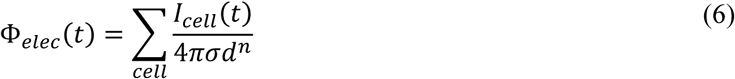

where *I_cell_* represents the current leaving the cell, *σ* is the medium conductivity and *d* is the distance between the center of the electrode and the cell. We used a *n* =1.6 to emulate the attenuation of spike amplitudes (Fig. S3a) due to realistic cell morphologies^55^.

#### Stimulation

During stimulation, electrodes were approximated as point current sources. The total injected current was estimated using the following Robin boundary condition^56^:

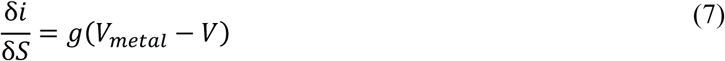

This simplification is possible because we used small electrodes, whose high impedances result in a nearly uniform current flow across the electrode surface (*V* ≪ *V_metal_*). Cell activation was induced by simulated synaptic events via NEURON’s NetCon/ExpSyn mechanism, following estimation of the steady-state value of transmembrane current ΔΦ*_m_*, which can be simplified to^57^:

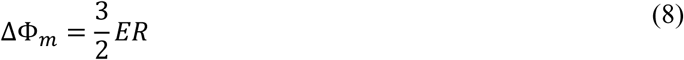

with:

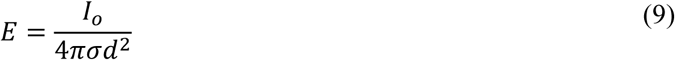

where *E* is the strength of the electric field, assuming a uniform conducting medium of infinite extent, *R* is the cell radius, *I_o_* is the current injected by the current source, and *d* is the distance from the current source. The corresponding synaptic conductance weight was then derived from the delivered charge. This stimulation model resulted in cell spike probabilities that are dependent on the distance to the electrode (Fig. S3b).

#### Spike detection

Extracellular spikes were identified through threshold detection, with the threshold set for each channel at 5 standard deviations (5σ) of the background noise RMS amplitude. A 3 ms refractory period was enforced to eliminate false positives from biphasic potentials.

### In vitro model

#### Cell cultures

The experiments followed both the European legislation regarding the use of animals for scientific purposes and the protocols approved by the ethical committee of i3S. The Animal Facility of i3S follows the FELASA guidelines and recommendations concerning laboratory animal welfare, complies with the European Guidelines (Directive 2010/63/EU) transposed to Portuguese legislation by Decreto-Lei no 113/2013, and is licensed by the Portuguese official veterinary department (DGAV, Ref 004461). Hippocampal tissues from embryonic (E18) Wistar Han rats were dissected in Hank’s Balanced Salt Solution (HBSS) and enzymatically digested in 1.5mg/ml trypsin (1:250) in HBSS without calcium and magnesium for 15 min at 37 °C. Trypsin was subsequently inactivated by adding HBSS supplemented with 10% foetal bovine serum and removed by washing. Tissue fragments were then mechanically dissociated using a plastic pipette, and the resulting cell suspension was filtered through a 40 µm cell strainer.

The dissociated neurons were cultured on six-well chamber MEA, 60-6well MEA200/30iR-Ti-rcr and MEA200/30iR-Ti-tcr (Multi Channel Systems, Germany), with a density of 4.5x10^4^ cells/well and 9x10^4^ cells/well, respectively. Cultures were maintained in Neurobasal^TM^ Plus medium (Thermo Fisher Scientific, USA) supplemented with 0.5 mM GlutaMAX, 2% (v/v) B27 Plus, and 1% (v/v) penicillin/streptomycin (P/S), with half-medium changes performed every 2-3 days. Each well of the six-well MEA has 9 titanium nitride (TiN) circular electrodes (array of 3x3), each spaced by 200 *μm* and with a diameter of 30 *μm*, plus a reference electrode. The experiments were performed between 17 and 42 days in vitro (DIV).

#### Electrophysiological recordings

We acquired extracellular signals using a MEA2100-Mini system (Multi Channel Systems, Germany) while maintaining cultures at 37°C and 5% CO_2_. We developed a custom C# application using the *McsUsbNet* library to interface with the MEA2100-Mini system. Signals were sampled at 10 kHz with a 0.1Hz hardware filter and processed with a 200 Hz high-pass Butterworth filter (2nd order). Spike detection was performed in real-time, with threshold crossings defined as peaks exceeding ±5× the RMS noise level. A 3 ms refractory period was enforced post-detection to prevent duplicate identification of biphasic waveforms. Stimulation artefacts were cleaned in real-time using the SALPA method^58^. Raw data and spike times were timestamped and stored for offline analysis.

#### Calcium imaging recordings

In calcium imaging preparations, neurons were transduced at 3 DIV with an adeno-associated virus (AAV) encoding for jGCaMP8m (ssAAV-DJ/2-hSyn1-jGCaMP8m-WPRE-SV40p(A); 6.6 × 1012 vg ml^−1^ titer^59^). As in previous studies^60^, the target viral load was of ten thousand viral particles per cell (i.e., multiplicity of infection (MOI) of 10k). The AAVs were produced by the Viral Vector Facility (VVF) of the Neuroscience Center Zurich (Switzerland). Concurrent MEA and calcium imaging experiments were performed at 18 DIV. Images were acquired by a sCMOS camera Prime 95B, 22 mm (Teledyne Photometrics, UK), mounted on a Nikon Eclipse Ti2-E (Nikon, Japan) inverted microscope with a Nikon Achro ADI 10X/0.25NA objective (with 1.5× optical zoom). Image acquisition was performed using Micromanager (Version 1.4) at 20 Hz (50 ms exposure). To synchronize MEA recording, stimulation, and image acquisition, the setup was triggered via a transistor-transistor logic (TTL) signal sent at the start of the protocol. Temperature (37 °C) and humidified 5% CO_2_ (stage top incubator, ibidi, Germany) were maintained via external controllers throughout the experiment.

#### Closed-loop control implementation

Both recording and closed-loop control operated on a dedicated workstation (Intel i7-6700, 32 GB RAM). Raw signals were streamed at 10 kHz via USB 3.0, with each batch containing synchronized samples across all electrodes. Additionally, the software enabled both precise MEA stimulator control via USB 3.0, implementing an automated closed-loop system with consistent <20 ms decision-to-stimulus latency. System reliability was maintained through error handling and continuous monitoring of data stream integrity, with validation tests confirming stable operation during extended experimental sessions.

### Problem Formulation

We formalized the task of optimal NB-evoking stimulation as a contextual multi-armed bandit problem^14^, where stimulation efficacy depends nonlinearly on both temporal and electrode context. Crucially, this nonlinear dependence operates within a partially observable system (the true network excitability state is obscured by measurement noise and unobserved latent variables) requiring policies that robustly handle stochastic outcomes.

#### State representation

The explicit design of a state representation to infer a hidden network state is well-precedented in adaptive stimulation^23,24^. We addressed this partial observability by approximating the neuronal population’s excitability level through a history of observed electrode activity. Each state *s_t_* at time step *t* was represented by a feature vector:

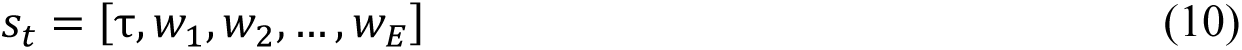

where τ is the elapsed time since last NB relative to the median of last five NIBI, *E* is the total number of electrodes, and *w*_(.)_ is the weight of a given electrode. The weight was calculated based on the number of spikes and their rank in individual NB for the last five NBs:

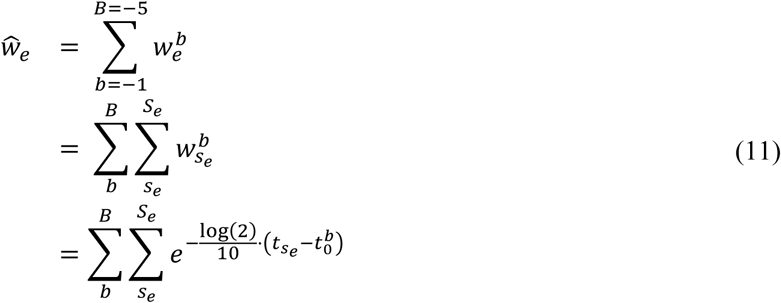

with *b* representing the index of the network burst, *s_e_* the index of the spike within electrode *e*, and 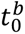 the starting time of network burst *b*. The individual electrode weights *w*_(.)_ were then calculated by normalizing *ŵ*_(.)_ with the sum of all the electrode weights. This representation is grounded on two key observations: (1) networks exhibit short-term stability in temporal dynamics (e.g., NIBI patterns), enabling predictive stimulation timing; (2) electrodes capturing earlier NB-onset activity likely reside closer to initiation sites, making them optimal stimulation targets for evoking subsequent NBs.

#### Action space

We defined an action as the delivery of a single monophasic cathodic pulse (−400 mV amplitude, 200 *μs* duration) through one electrode (or none). The pulse parameters were optimized for sparse neuronal networks, where −400 mV serves as a near-threshold stimulus capable of evoking NBs while maintaining localized activation. Higher amplitudes (e.g., −800 mV) were avoided as they produce broader activation radii, despite remaining within safe charge injection limits. These parameters also minimize Faradaic processes at the electrode interface and preserve energy efficiency.

#### Step length

The step length critically shapes both the temporal resolution of evoked actions and the informational richness of the state. A 200 ms inter-step interval was chosen to (1) allow full recovery of neuronal membranes after each stimulation, preventing cumulative depolarization blocks; and (2) accommodate typical cell refractory periods (1–10 ms) and network-level response latencies (tens to hundreds of milliseconds) observed in the cultures. This interval ensured minimal interference from preceding pulses while enabling observation of evoked network activity.

#### Episode initiation and termination

Because the task required evoking a network burst (NB), each episode was terminated upon detection of a NB onset. To account for natural variability in NB duration (which could depend on factors like network maturation or time since medium replacement), the subsequent episode was initiated only after confirming the end of the previous NB. This was implemented by detecting a minimal quiescence period of 200 ms following each NB, ensuring network readiness for the next stimulation cycle.

#### Reward function

An immediate reward *r_t_* = *R*(*s_t_*, *a_t_*, *s_t_*_+1_) is received by the RL agent after taking action *a_t_* at *s_t_* and observing new state *s_t_*_+1_. The reward function was designed to promote evoked network burst (NB) and correct waits (no stimulation), and penalize failed stimuli and spontaneous NB:

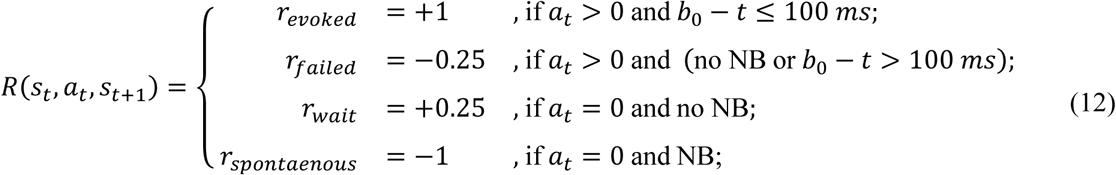

#### Actor-Critic framework

We adapted Proximal Policy Optimization^32^ (PPO) by modifying the critic to estimate *Q*(*s*, *a*) rather than *V*(*s*) (Supp. Algorithm 1). This modification was motivated by our contextual bandit framework, where stimulation decisions are treated as independent and sparse NB evocations require precise credit assignment. Whereas a state-value critic would conflate the effects of all electrode choices, our Q-value critic directly models action-specific outcomes, enabling clearer identification of which actions resulted in successful NB evocation. We used batched training with 256 samples per batch, performing policy updates over 10 epochs using 64-sample mini-batches with advantage normalization to improve learning stability. The architecture consisted of single 32-unit layer MLPs for both the actor (policy) and critic (Q-value) networks, with the policy network outputting a categorical distribution (*softmax* activation) over the possible actions.

PPO training used a discount factor of γ = 0 (consistent with the myopic nature of the contextual bandit formulation), a learning rate of 3 × 10^-4^, an entropy coefficient of 0.001, and a clipping threshold of 0.2.

#### Convergence criteria

To quantitatively determine when an online-trained agent stabilized upon an effective control policy, we established a dual-condition convergence threshold based on episodic cumulative reward. Raw cumulative rewards were first smoothed using a centred rolling median filter (window size = 30 episodes). Policy convergence was defined as the first episode where the trailing window of median-smoothed rewards simultaneously satisfied two criteria: (1) absolute efficacy, requiring all values within the window to exceed a minimum reward threshold of 1.5 (which is roughly equivalent to waiting at least two steps more than the number of stimulations applied, and successfully eliciting a NB); and (2) behavioural stability, requiring the standard deviation of the rewards within the window to remain below a strict tolerance of 0.2. The exact time and step count to reach this conservative convergence point were recorded to evaluate learning speed. Networks that failed to satisfy both criteria prior to the termination of the training phase were labelled as non-converging.

#### Policy distillation

We implemented a policy distillation^61^ approach that transfers knowledge from multiple network-specific policies into a unified *Generalist* policy. The distillation process was formulated as a supervised learning task using an aggregated dataset of state-action pairs collected from the *Specialist* agents (128 samples per agent). The *Generalist* agent (a 32-unit single layer MLP) was trained to minimize the Kullback-Leibler (KL) divergence between its predicted action distribution *π_Generalist_*(*a*|*s*) and the action distributions of *N Specialist* policies 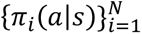. This objective was optimized using Adam (learning rate = 0.001) with batched updates (512 samples per batch) drawn from this stratified dataset.

### Experimental protocols

#### General procedures

All *in vitro* validation experiments followed a standardized temporal structure to ensure comparability. Each recording session began with a 3-minute initial baseline to assess spontaneous activity and concluded with a 3-minute final baseline to evaluate network stability. Distinct policy phases (e.g., *Generalist*, *Specialist*, and Controls) were separated by 30-second no-stimulation intervals to minimize immediate carry-over effects. To ensure immediate agent responsiveness, we pre-loaded the state buffer with a sequence of five 1.8 s intervals. This prevented the agent from delaying stimulation onset in networks with long spontaneous NIBIs.

#### Protocol 1: Transfer vs. Online Learning (Generalist vs. Specialist)

To benchmark the performance of the pre-trained *Generalist* policy against the online learning capabilities of the *Specialist* policy, we implemented a counterbalanced crossover design. Following the initial baseline, the *Generalist* policy (1 minute) and the *Specialist* policy (10 minutes) were applied in one of two testing sequences (S→G or G→S) to control for potential order effects.

#### Protocol 2: Generalist vs. Heuristic Controls

To dissect the efficacy of the *Generalist* policy, we compared it against two heuristic controls: *Without Best* (which excluded the single highest-performing electrode identified during the *Generalist* phase) and *Random* (which maintained the *Generalist* policy’s stimulation usage but randomized stimulation timing and electrode selection). Following the initial baseline, the *Generalist* policy (1 minute) was applied first to establish the target metrics. This was followed by the two control policies (1 minute each), applied in a randomized order. This sequential design ensured that control policies were conditioned on the specific active electrodes identified by the *Generalist* agent. The *Without Best* policy specifically tested the contribution of high-performing electrodes, whereas the *Random* policy served as a stochastic control, together providing complementary insights into policy-dependent network responses.

### Signal processing

#### Network burst detection

Network bursts (NBs) are transient periods of coordinated spiking activity across the electrodes and were defined by three criteria: (1) the occurrence of successive spikes across any electrodes within ≤20 ms (global inter-spike interval threshold); (2) the participation of at least 3 electrodes (1/3 of total array coverage), each contributing with ≥3 spikes. (3) NBs occurring within a brief integration window (200 ms) are merged into sustained events, accounting for potential spatial propagation and reverberatory activity (sometimes called “superbursts”). Unlike conventional binning methods^62^, this method provided exact burst onset timing (i.e., to the first spike), which is essential for our state representation (see *Methods: State representation*).

#### Metrics of network activity

We systematically quantified neuronal network dynamics using five complementary metrics designed to capture distinct functional aspects. The network burst rate (NBR) measured the frequency of coordinated bursting events per minute, reflecting overall network excitability. The network burst duration (NBD) quantified the average interval between burst onset and offset, with longer durations indicating enhanced network-level sustained firing capability. Bursting patterns were further characterized through the network inter-burst interval (NIBI, the mean time between consecutive burst offsets and onsets) and its coefficient of variation (NIBI_CV_), which together described the temporal organization of network bursts. Finally, we assessed network synchronization using an adapted *χ* metric^63^ that compares the variance of population-wide activity to mean single-electrode variances. To ensure analysis of functionally relevant channels, we excluded electrodes with insufficient firing rates (<0.1 Hz global average), effectively filtering out quiescent or poorly coupled recording sites.

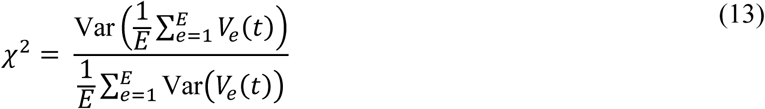

where *E* is the number of electrodes, and *V_e_*(*t*) corresponds to the binarized activity of each electrode, obtained by convolving spike trains with a 50 ms square wave kernel. This approach specifically detects coordinated activation patterns across electrodes while remaining insensitive to differences in spike counts per activation event, a critical feature given potential variability in local neuron densities surrounding each electrode.

#### Fluorescence analysis

To extract neuronal activity traces, individual cell bodies were manually identified by defining regions of interest (ROIs) for each perceptible soma using ImageJ (version 2.16.0). Raw fluorescence intensity was then extracted from these ROIs and converted to relative fluorescence changes (*ΔF*/*F*) to account for background noise and correct for slow baseline drifts. For each frame, the normalized signal was calculated using the following equation:

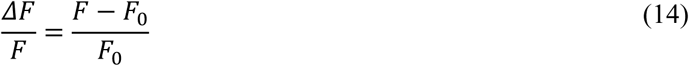

where *F* represents the fluorescence intensity of the current frame and *F_0_* is the baseline fluorescence, defined as the median intensity within a one-second sliding window centred on the frame.

### Statistics and reproducibility

Given that most studied variables exhibited non-normal distributions, data are presented as the median with interquartile ranges, or with specific percentile boundaries (e.g., 25th–75th or 10th–90th percentiles) as explicitly defined in the figure legends. For comparisons between two independent groups (e.g., *in silico* versus *in vitro*, or different policy orders), the two-sided Mann-Whitney U test was used. For comparisons of the same network or seed under different policies, two-sided paired permutation tests were used for two conditions, and the Friedman test followed by post-hoc paired permutation tests for three or more conditions. We employed the Kendall correlation to test potential relationships between network excitability and policy performance. Multiple comparisons were adjusted using the Holm-Bonferroni correction.

Sample sizes (*N*) are reported as exact values throughout the text and figure legends. For *in silico* experiments, *N* represents independent network instantiations with different seeds. For *in vitro* experiments, it represents individual MEA recordings. These recordings were derived from 132 separate E18 rat culture wells across 15 independent biological preparations. Only wells exhibiting spontaneous bursting activity were considered for experimentation, as model validation and policy evaluation require bursting as a precondition. Because network activity profiles change significantly over the course of maturation, some wells were recorded up to three times at widely separated days in vitro (spanning DIV 17–42) to capture the full phenotypic variance of the developing networks. These distinct temporal states were treated as individual observations for the overall distribution.

## Data Availability

The data that support the findings of this study are openly available in Zenodo at [link to be made public after manuscript is accepted for peer review]

## Code Availability

Code for 1) the neuronal population model developed in NEURON simulation environment and recreating in vitro MEA recordings/stimulation, and 2) the C# application for real-time control of the MCS-MultiChannel Systems MEA2100 electrophysiology hardware, are freely available for download in GitHub at [repository and link to be made public after manuscript is accepted for peer review]. All data, models and code are openly available and in accordance with FAIR data principles.

## Author Contributions

E.C. and P.A. conceived and designed the study. E.C. developed the *in silico* model and carried out all *in silico* experiments, electrophysiological recordings, data analysis, and statistical analysis. J.M. and M.A. were responsible for culture preparation and maintenance. Calcium imaging recordings were performed jointly by E.C. and J.M., and the subsequent analysis was conducted by E.C. and R.P. E.C. and P.A. wrote the manuscript. All authors contributed to the interpretation of the results, critical revision, and final approval of the manuscript. P.A. provided overall supervision of the study.

## Acknowledgements

This work was supported by Foundation ‘la Caixa’ - CaixaResearch Health 2022 (grant HR22-00189). E.C. was supported by Fundação para a Ciência e a Tecnologia (FCT, grant 2024.03120.BD) in the scope of the BiotechHealth PhD Program (Doctoral Program on Cellular and Molecular Biotechnology Applied to Health Sciences). The authors thank all the members of the Neuroengineering and Computational Neuroscience group for help and critical discussions.

## Ethics declarations

The experiments followed both the European legislation regarding the use of animals for scientific purposes and the protocols approved by the ethical committee of i3S. The Animal Facility of i3S follows the FELASA guidelines and recommendations concerning laboratory animal welfare, complies with the European Guidelines (Directive 2010/63/EU) transposed to Portuguese legislation by Decreto-Lei no 113/2013, and is licensed by the Portuguese official veterinary department (DGAV, Ref 004461).

