## Supplementary material for "Closed-loop control of *in vitro* neuronal activity using reinforcement learning after in silico pre-training": Figure S# / Table S#

### Supplementary Information

**Figure S1.** Validation of performance stability across consecutive *Generalist* policy phases

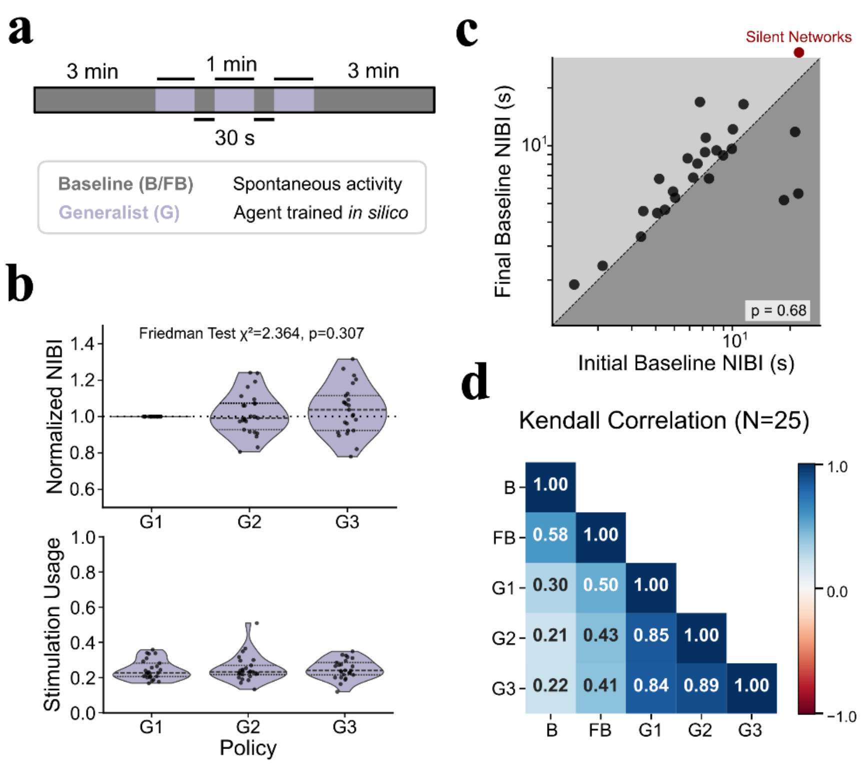

**(a)** Schematic of the experimental protocol. Same structure as Protocol 2 from main text, but with three *Generalist* phases (G1, G2, G3) separated by 30 s inter-phase intervals. **(b)** Top: Normalized NIBI across the three Generalist phases (normalized to G1); the Friedman test indicates no significant difference. Bottom: Stimulation usage across phases remains identical. **(c)** Quantification of network fatigue. Scatter plot of spontaneous NIBI before (Initial Baseline) and after (Final Baseline) policy deployment. While most networks showed a slight tendency toward increased NIBI, three networks became markedly more bursty, and one network was silenced entirely. **(d)** Kendall rank correlation matrix of NIBI across all phases (N=25). Moderate correlation between initial (B) and final baselines (FB) indicates temporal stability of intrinsic network dynamics, while high correlations between baseline and active stimulation policies (G1, G2, G3) confirm preserved network ranking across consecutive policy deployments.

**Figure S2.** Functional exceptions to somatic proximity as a predictor of spiking activity and electrode efficacy

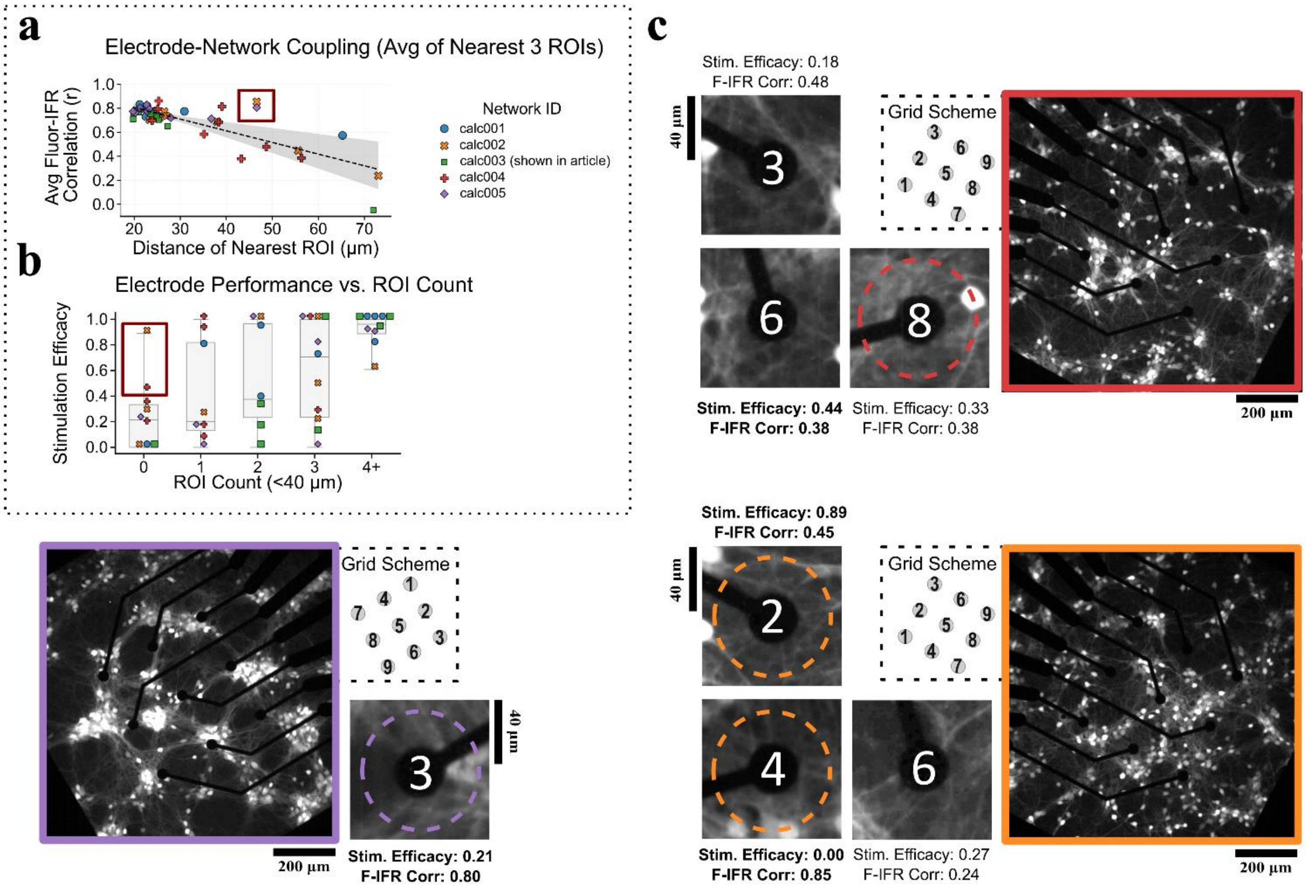

**(a)** Scatter plot showing the relationship between the distance to the nearest ROI and the average fluorescence–IFR correlation across all electrodes (N = 45, 5 cultures with 9 electrodes each). Correlation tends to decay beyond ~40 µm, as captured by the linear trend (dashed line) and 95% CI (shaded area). The red box highlights two electrodes that maintain high correlations despite lacking visible ROIs within 40 µm, further examined in **c**. **(b)** Stimulation efficacy as a function of local ROI count (< 40 µm) across all electrodes (N = 45). The red box highlights electrodes with zero adjacent ROIs yet efficacy above 0.40 (8/9 and 4/9 recordings, respectively), further examined in **c**. Low but non-zero efficacy values should be interpreted cautiously, as network bursts occurring spontaneously after stimulation may inflate these estimates. **(c)** Calcium imaging fluorescence frames of the three networks (border colors match scatter plot markers) containing zoom-ins and statistics of some 0-ROI electrodes, including the highlighted exceptions in **a** and **b** (in bold). Dashed circles indicate the 40 µm search radius. Each zoom-in displays the electrode number (matching the grid scheme, inset), stimulation efficacy, and fluorescence–IFR correlation. Purple: electrode 3 shows high network correlation (0.80) and adjacent somata partially hidden by the opaque metallic lead, that become noticeable with increased brilliance. Red: electrode 6 show moderate efficacy (0.44) and low correlation (0.38) with no visible adjacent ROIs. Orange: electrode 4 shows high correlation (0.85) with no visible adjacent ROIs; electrode 2 shows high efficacy (0.89) with no visible adjacent ROIs, suggesting distal network coupling as an alternative basis for excitation. Neuronal processes are visible across all zoom-ins.

**Figure S3.** Distance-dependent activation and signal detectability in simulated neurons

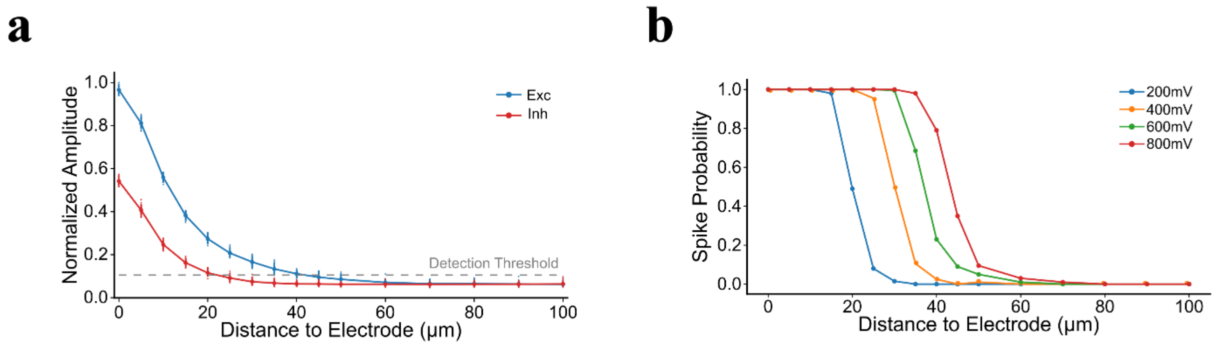

**(a)** Normalized amplitude of detected spikes from excitatory (blue) and inhibitory (red) neurons as a function of distance to the virtual electrode. The dashed line indicates the detection threshold (5σ of the background noise RMS); excitatory spikes fall below it at approximately 40 µm, while inhibitory spikes, starting at a lower amplitude, do so at a shorter range. **(b)** Spike probability as a function of distance to the virtual electrode for four stimulation amplitudes (200–800 mV). Higher amplitudes extend the effective activation range, with the 50% spike probability threshold shifting from ~20 µm at 200 mV to ~50 µm at 800 mV.

Table 1. Overview of the parameters of the neuronal models.

| Parameter | Description | Unit | Pyramidal | Interneuron |
|---|---|---|---|---|
| $d$ | soma diameter | $\mu m$ | 20 | 15 |
| $C_m$ | membrane capacitance | $\mu F \cdot cm^{-2}$ | 1 | 1 |
| $g_K$ | maximum potassium conductance | $mS \cdot cm^{-2}$ | 36 | 20 |
| $g_{Na}$ | maximum sodium conductance | $mS \cdot cm^{-2}$ | 120 | 80 |
| $g_l$ | leak conductance | $mS \cdot cm^{-2}$ | 0.1 | 0.042 |
| $E_K$ | Nernst potential of potassium | $mV$ | -80 | -80 |
| $E_{Na}$ | Nernst potential of sodium | $mV$ | 70 | 70 |
| $E_l$ | Nernst potential of the leak current | $mV$ | -65 | -65 |
| $V_T$ | potential to adapt spike threshold | $mV$ | -55 | -55 |
| $g_{noise}$ | maximum noise conductance | $nS$ | 0.15 | - |
| $\tau_{noise}$ | noise time constant | $ms$ | 3 | - |
| $r_{noise}$ | noise mean firing rate | $Hz$ | 200-300 | - |

Table 2. Synaptic parameters.

| **Source** | **Target** | **Receptor** | $\tau_{rise}$ **(ms)** | $\tau_{decay}$ **(ms)** | $\hat{g}$ **(nS)** | **U1** | $\tau_F$ **(ms)** | $\tau_D$ **(ms)** |
|---|---|---|---|---|---|---|---|---|
| Pyr | Pyr | AMPA | 0.1 | 3.0 | 0.58 | 0.5 | 17.0 | 671.0 |
| | | NMDA | 3.9 | 75-125 | 0.1 | 1 | 0.1 | 671.0 |
| Pyr | PV+ | AMPA | 0.1 | 4.12 | 2.0 | 0.23 | 10.0 | 410.0 |
| PV+ | Pyr | $GABA_A$ | 0.1 | 5.94 | 2.15 | 0.16 | 8.6 | 965.0 |

#### Algorithm 1. Adapted Proximal Policy Optimization with Q-Critic

**Require:** $\pi_\theta$ (actor), $Q_\omega$ (critic), $\alpha_\theta$, $\alpha_\omega$ (learning rates), $\beta$ (entropy weight), $N$ (batch size) = 256, $M$ (mini-batch size) = 64, $K$ (epochs) = 10

**Ensure**: Actor and critic jointly optimized for sparse, action-specific rewards

Initialize parameters $\theta$, $\omega$

Initialize a storage buffer$\mathcal{D} = S, A, R$

**for** each episode **do**

  Get initial state $s_0$

  **while** no network burst **do**

    Sample action $a_t \sim \pi_\theta(a_t | s_t)$

    Execute $a_t$, observe reward $r_t$ and next state $s_{t+1}$

    Store transition $(s_t, a_t, r_t) \in \mathcal{D}$

    **if** $|\mathcal{D}| \ = N$ **do**

      Compute targets $y_t = r_t$ (contextual bandit, $\gamma = 0$)

      Compute advantages $A_t = y_t - \sum_{a'} \pi_\theta(a' | s_t) Q_\omega(s_t, a_t)$

      Cache log $\pi_\theta^{old}(a_t | s_t)$

      **for** epoch $k = 1$ to $K$ **do**

        Randomly sample mini-batch of $M$ transitions from $\mathcal{D}$

        Compute updated log-probs log$\pi_\theta(a_t | s_t)$

        Compute ratios $\rho_t = \exp\left(\log\pi_\theta(a_t | s_t) - \log \pi_\theta^{old}(a_t | s_t)\right)$

        Compute entropy $\mathcal{H}_t = -\sum_a \pi_\theta(a_t | s_t) \log \pi_\theta(a_t | s_t)$

        Normalize advantages $\hat{A}_t = \frac{A_t - \mu}{\sigma}$, with $\mu$, $\sigma$ computed within the mini-batch

        Compute actor loss $L_\theta = -\frac{1}{M} \sum_{t=1}^{M} \min\left(\rho_t \hat{A}_t, \text{clip}(1 - \rho_t, 1 + \rho_t) \hat{A}_t\right) - \beta \mathcal{H}_t$

        Compute critic loss $L_\omega = \frac{1}{M} \sum_{t=1}^{M} \left(y_t - Q_\omega(s_t, a_t)\right)^2$

        Update $\theta \leftarrow \theta - \alpha_\theta \nabla_\theta L_\theta$

        Update $\omega \leftarrow \omega - \alpha_\omega \nabla_\omega L_\omega$

      **end for**

      Clear buffer $\mathcal{D} \leftarrow \emptyset$

    **end if**

  **end while**

**end for**
